# Long-term population monitoring reveals an alarming decline in the endangered endemic Réunion Harrier *Circus maillardi*

**DOI:** 10.1101/2025.11.07.687144

**Authors:** Alexandre Villers, Thomas Cornulier, Cyril Eraud, Thibaut Couturier, Pierrick Ferret, Damien Chiron, Nicolas Laurent, Aurélien Besnard, Vincent Bretagnolle, Marc Salamolard, Valérie Grondin, François-Xavier Couzi, Rémi Fay, Steve Augiron

## Abstract

Insular species are disproportionately vulnerable to extinction due to small population sizes, geographic isolation, and high susceptibility to stochastic and anthropogenic pressures. The Réunion Harrier (*Circus maillardi*), the last breeding raptor on La Réunion (Indian Ocean), is currently listed as Endangered (EN) on the IUCN Red List, although population decline has so far only been suspected rather than robustly quantified. Concerns have recently emerged regarding the demographic impacts of secondary poisoning from anticoagulant rodenticides used for rat control in agricultural landscapes, in addition to habitat loss, fragmentation, and reduced genetic diversity. Here, we analysed data from three standardised island-wide breeding surveys (1998–2000, 2009–2010, 2017–2019), encompassing 346 sampling points and totalling over 1,500 h of observation, to quantify population trends and their spatial variation. Using generalized additive mixed models accounting for spatial structure and heterogeneous survey designs, we detected a significant decline of -46% [-62%; -27%] in the relative abundance of breeding pairs at the island scale over 21 years, equivalent to three generations, supporting the species’ current Endangered status. The decline was steeper between 1998 and 2009 (-33% [-52%; -9%]) than between 2009 and 2017 (-18% [-42%;+14%]). The decrease was more markedly pronounced in the eastern part of the island (-57% [-71%; -40%]) than in the west (-29% [-53%; + 0.8%]), reflecting a complete reversal of the historical east-west abundance gradient. Combined with previous evidence of inbreeding, high mutational load, and rodenticide exposure, these results suggest the population is undergoing an extinction vortex requiring immediate, spatially targeted conservation actions, including reduced rodenticide exposure and genetic management. These findings also underscore the need for sustained monitoring and improved demographic knowledge to support the implementation of the ongoing National Species Action Plan.

## INTRODUCTION

The current biodiversity crisis has led to the extinction of hundreds of species during the past 500 years, and hundreds more are on the verge of extinction (Ceballos *et al*., 2017; Ceballos & Ehrlich, 2023). While this crisis is affecting all taxa across the globe (see e.g. Alroy, 2015; Régnier *et al*., 2015; McClure et al., 2018), some geographical areas such as islands are more sensitive to extinction risk (Wood *et al*., 2017). As a whole, islands account for 61% of known extinctions and 37% of critically endangered species worldwide (Tershy *et al*., 2015). Indeed, insular communities are often characterised by low population sizes and geographical isolation, both increasing not only the sensitivity of those populations to environmental stochasticity (Lande, 1998) but also exposing them to genetic drift (Funk *et al*., 2016). The latter can result in inbreeding, increasing the emergence and persistence of deleterious mutations (Keller & Waller, 2002). All these mechanisms can have detrimental effects on the viability of insular animal populations, which can then be reinforced by anthropogenic pressures, such as the introduction of invasive species (Blackburn *et al*., 2014), habitat modification (Gillespie *et al*., 2008) or recreational activities(Doherty *et al*., 2021).

La Réunion, a 2,500 *km*^2^ island located in the Indian Ocean, belongs to the Madagascar biodiversity hotspot, one of the 25 areas of biodiversity importance identified by Myers *et al*. (2000), characterized by a high level of endemism. However, since humans settled on the island, in the middle of the XVII^th^, habitat loss, fragmentation and modification have been unusually rapid (Strasberg *et al*., 2005). This includes the introduction of numerous invasive plant and animal species (Baret *et al*., 2006), with harmful effects to native fauna and flora. The most notable impacts stem from predation by feral cats *Felis catus* and black rats *Rattus rattus*, or competition and habitat modification by invasive plants such as *Miconia calvescens*. Thus, during the past 300 years, 73% of the native vegetation was transformed (Lagabrielle *et al*., 2009), and 51% of known native bird species were extirpated from the island (Mourer-Chauviré *et al*., 2006). Today, nine endemic bird species are still present on the island, the Réunion Harrier *Circus maillardi* being the only breeding bird of prey remaining on La Réunion (Safford & Hawkins, 2020).

This raptor faces threats similar to those affecting raptors worldwide (O’Bryan *et al*., 2022), such as direct destruction or trapping (Thomason *et al*., 2023), collisions with power lines, wind turbines (Bellebaum *et al*., 2013; Slater *et al*., 2020) or vehicles (Bullock *et al*., 2024). In addition, the species is subject to poaching, for local folklore. Secondary poisoning from anticoagulant rodenticides (hereafter ARs, Coeurdassier *et al*., 2019), used to control rat populations in farmlands, is considered one of the most severe threats (BirdLife International, 2021), given that rodents constitute a large proportion of harriers’ diet. Recent genetic analyses also highlighted a high degree of inbreeding which may result in compromised genetic health and reduced adaptive potential (Bourgeois *et al*., 2024). In this context, Fay *et al*. (2023) developed a demographic model of the species’ dynamics which suggested that the population was declining at an alarming rate. Their results underlined a risk of quasi-extinction in the near future (c. 40 years). However, this model relied on fragmentary demographic data, mostly from the eastern part of the island, and therefore the predicted population trajectory requires validation, to precisely quantify ongoing population changes. The absence of reliable data on recent population trends has hindered the objective reassessment of the species’ Red List status. Currently, the IUCN endangered (EN) status of the Réunion Harrier relies on criteria related to its geographic range and habitat (criteria B1ab(iii)), as well as on a population size estimated to number fewer than 250 mature individuals (criterion D), whereas the decline itself is only suspected (BirdLife International, 2021).

In 2022, a national conservation action plan (2022-2031) was launched to improve the species’ conservation status by reducing identified threats. The plan calls for these actions to rest on sound knowledge of the species’ ecology and population dynamics. However, the surveys conducted over the past 25 years differed in sampling schemes and protocols, so raw counts cannot support reliable inference on population trends. Here, we assess changes in the abundance of the Réunion Harrier breeding population over the past 20 years, using three island-wide surveys (1998-2000, 2009-2010, and 2017-2019). Considering results of a genetic studies (Bourgeois *et al*., 2024) and the high exposures to anticoagulant rodenticides, which are mainly used in human-modified environments, (Coeurdassier *et al*., 2019), we suspected geographical contrasts in demographic functioning. We therefore modelled spatial structure in population trends explicitly, which may help elucidating the underlying demographic processes at play, and target conservation actions spatially.

## MATERIAL & METHODS

### Study species

The Réunion Harrier is a medium-sized raptor (700 – 1,000g), formerly present on Mauritius but going extinct soon after the arrival of the first European settlers (Cheke, 2010). It is closely related to the Malagasy Harrier *Circus macrosceles*, that breeds on Madagascar, from which it diverged c.a. 0.1-0.3 mya ago (0.05 - 0.5 CI, see Oatley *et al*., 2015). On La Réunion, the species can be encountered throughout the island, with most breeding pairs located between the coast and 1,200m (Augiron, 2022). It originally exploited native forests (Clouet, 1978) that covered most of the island but has adapted to the degradation and modification of native habitats by humans. For example, it now breeds in the low vegetation of secondary forests and preys upon introduced mammals, which have proliferated significantly in the most heavily modified habitats. The black rat in particular thrives in agricultural landscapes, especially in sugar cane fields and near cattle farms, and thus constitutes a significant portion of the harrier’s diet (Augiron, 2022). The most recent survey suggests a population size of approximately 150 breeding pairs, and approximately 560 mature individuals in 2009-2010 (Grondin & Philippe, 2011), whereas Ghestemme et al. (1998) reported a population of 120-180 breeding pairs.

### Data collection

Three surveys were conducted over the past two decades: in 1998-2000 (Bretagnolle *et al*., 2000; Ghestemme et al., 1998), 2009-2010 (Grondin & Philippe, 2011) and 2017-2019 (Chiron & Augiron, 2019), hereafter referred to as 1998, 2009 and 2017 surveys. All surveys shared a common field methodology, based on counts of individuals on a set of sampling points (hereafter referred to as “SP”) distributed over the island in different habitats (see Fig. 1).

**Figure 1:**
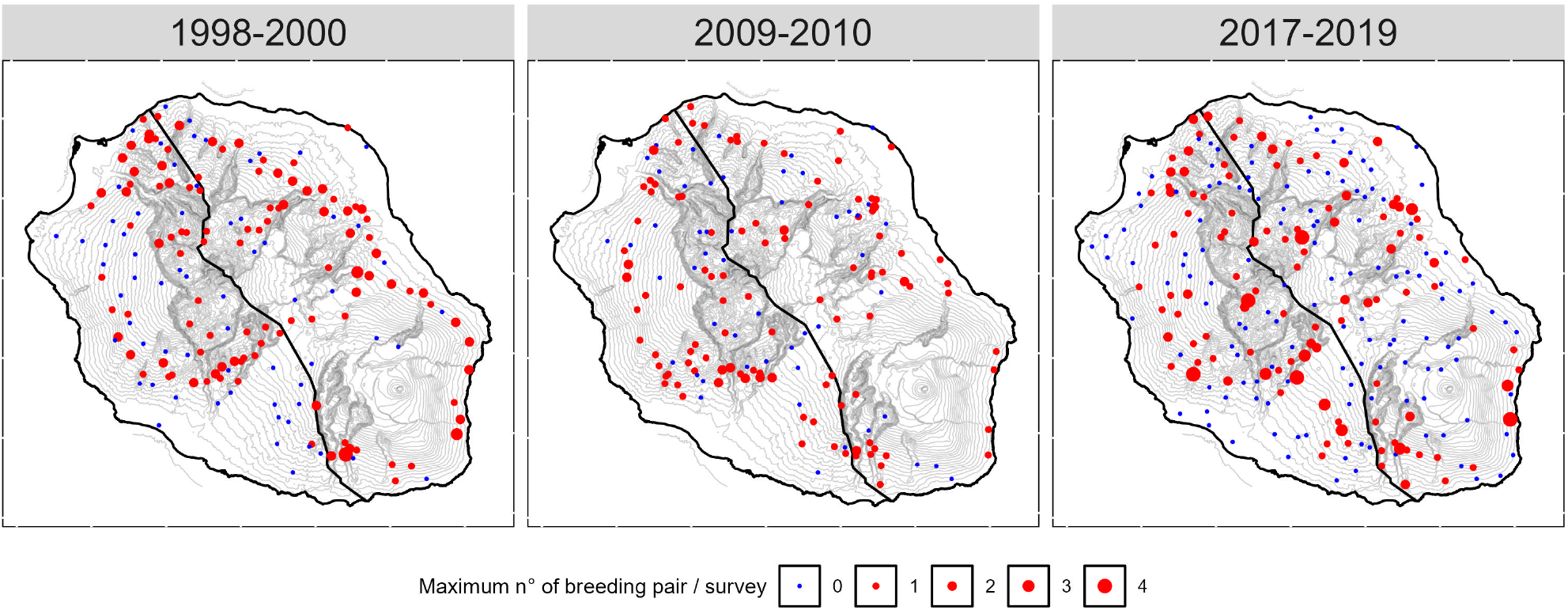
Locations of observation points for the three different surveys (1998-2000, 2009-2010, 2017-2019) on La Réunion. The colour of the points indicates whether a breeding pair was observed (red) or not (blue). Point size is proportional to the maximum number of pairs detected at each sampling point during each survey. Polygons delineate the eastern and western areas used for trend comparisons. Contour lines represent altitude isopleths every 100 meters, computed from the BD ALTI 2.0 (©IGN)

SPs were located near paths, roads, forest tracks or mountain trails to facilitate to facilitate access, and on elevated vantage points to ensure a wide field of view (Couturier *et al*., 2022). Over the three surveys, 346 unique SPs were monitored at least once, with 37.5% (n=133) and 15.5% (n=55) of those points that were common to two and three surveys, respectively. On each SP, one or repeated observation sessions (hereafter session) were performed. The total number of sessions was 341 in 1998, 185 in 2009, and 522 in 2017. Each SP was, on average, sampled three times (range: 1-8) over the three survey periods. Survey conducted in 2017 had a more extensive coverage of the island, and notably embraced a wider altitudinal gradient than previous surveys. Average altitude was 798 m (range: 2m – 2482 m) in 1998, 756 m (range: 2m – 2327m) in 2009 and 856 m (range: 2m – 2482m) during the last survey. In 2017, if the visual range on a SP, previously surveyed in 1998 and/or 2009, had been obstructed by vegetation or a newly constructed building, the observer was allowed to relocate the SP within a 200 m radius of the original location, and the identity of the SP was preserved over the 3 surveys. Otherwise, a new identity was attributed to the SP. Sessions on each SP started on average 217 *±* 103 min. (SD) after sunrise 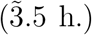. All observations were conducted only during favourable weather conditions (no or low wind and no rainfall or very light showers). If the weather conditions were detrimental to the activity of the harriers or to their detection (fog, heavy showers, strong winds), the session was interrupted, and interruptions were subtracted so that only effective observation time was counted. In case the conditions did not improve, the session was cancelled, and a new one was scheduled in the following couple of weeks. Planned session length was 120 min in 1998 and 2009, and 80 min in 2017. Overall, the duration of observations ranged from 25 to 300 min., with an average duration of 112 *±* 26 min. in 1998, 114 *±* 26 min. in 2009 and 80 *±* 0 min. in 2017. During the first two surveys, the number of observers was limited, and all participants were professionals from conservation organizations. For the third survey, a citizen science approach was implemented to significantly increase the number of sampling points (Couturier *et al*., 2022). Except for SEOR’s staff members or experienced ornithologists, all participants received both theoretical and practical training before their participation. This mandatory training helped to standardize ornithological skills (such as identifying sex, age, and the various reproductive behaviours) and ensured that participants understood the objectives and the protocol. Breeding activity was known to peak between April and late June only from 2010 onwards (Grondin & Philippe, 2011), so this information was unavailable for the first survey. Consequently, in 1998, observations spanned the whole year, with only 33% of samples collected between April and June, whereas the 2009 and 2017 surveys targeted this period (95% and 84% of samples, respectively, see also Fig. S1).

During each session, all birds that could be detected with the naked eye from the SP were reported on a field map (Couturier *et al*., 2022). Binoculars were then used to determine sex and age, and to look for distinctive features (plumage coloration, moult, wing tags) that could help distinguishing individuals in later observations. For all surveys, if one or more individuals exhibited reproductive behaviour, the observer reported it along with the time and location of the event, the sex and age of the birds concerned (according to the possible combinations proposed for each behaviour see Tab. 1), and noted the presence of distinctive signs if applicable. The observer then assigned an identifier to the individual or pairs. When the same individuals or pairs were re-observed during one observation session, the same identifier was used to keep track of individual information. To remain conservative, an individual or pair was counted as new only if i) wing tags or plumage features (e.g. colour variation, missing feathers, ongoing moult) showed it to be a different individual, or reproductive behaviour occurred in an area without prior observation; and ii) individuals were observed simultaneously, or within a very short interval, at locations too far apart for double counting. Observers recorded every individual or pair showing at least one reproductive behaviour and classified the degree of breeding evidence using the criteria in Tab. 1. At the end of each session, the observer estimated the minimum number of confirmed, probable and possible pairs. Only the “probable” and “confirmed” categories were used in our analysis, since the category “possible” was not available for the first survey.

**Table 1:** Standardised criteria applied across all surveys to classify the degree of breeding evidence. For the analyses, only those observations categorised as “confirmed” or “probable” breeding were retained.

| **Behaviours** | **Status** | **Description** |
|---|---|---|
| Prey transfer | **Confirmed** | Young following an adult carrying a prey and/or prey transfer between adults |
| Carrying a prey |  | Adult carrying a prey for young at nest |
| Young’s |  | Young with persistent begging calls |
| Nest material | **Probable** | Adult bringing nest material |
| Copulation |  | Observation of a copulation |
| Nuptial displays (2 adults) |  | Pair that performed nuptial displays |
| Pair displayed together |  | Two adults of opposite sexes that displayed together |
| Alarm call |  | Alarm call of an adult near a potential nest site |
| Adult landing frequently | **Possible** | Adult repeatedly landing in the same place |
| Solicitation |  | Adult females that gave solicitation to passing adult male |
| Territorial defence |  | Aggression towards another individual, territorial display |
| Nuptial displays (1 adult) |  | Single adult that performed nuptial displays |

### Statistical analysis

To estimate breeding population trends of La Réunion Harrier reliably, we had to account for changes in sampling effort across surveys, which differed in both spatial (sampled locations changed) and temporal coverages. Some points were indeed sampled in only one survey, others repeatedly. We did so using Generalized Additive Mixed Models (Wood, 2017, package *mgcv 1*.*9-3* ), which handle non-linear relationships, and spatial and temporal dependencies as well. This approach was particularly suited to the data set generated by the three surveys, with varying sampling effort. The response variable was the number of breeding pairs observed (*ptot*, i.e. pairs recorded as probable or confirmed breeders, cf. supra) at each point and each observation session, interpreted here as a relative abundance index. This response variable was assumed to follow a Tweedie distribution (see S2 for details). The effects of explanatory variables were modelled with splines for *Date*, log observation *Effort*, coordinates of sampling points (*X, Y* ). Year and site identity, expressed as factors *Y earF* and *SiteID*, were included as random intercepts to account for between-year and between-site variation:

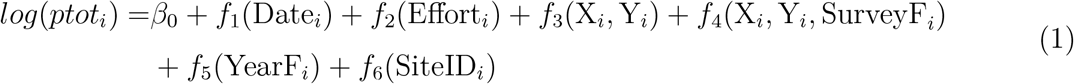

where *f*_*j*_ is a smooth function of covariates (Wood, 2017).

To assess whether the model adequately captured the spatio-temporal heterogeneity of sampling (at a reviewer’s request), we fitted a simpler model to four datasets: the full dataset; only points common to the three surveys; only points surveyed between April and June; and points common to the three surveys and surveyed between April and June. Because trends were of similar magnitude across datasets (see Figure S5), we concluded that the main model handles this heterogeneity adequately and present only its results.

To compute relative population trends, we used model predictions per sampling point, conditional on survey and location, and other covariates fixed at their respective mean value. This can be interpreted as a population index proportional to density, with an arbitrary scale. Confidence intervals were obtained by posterior simulation, following (Wood, 2017, chapter 6.10, p. 293). Details on the approach can also be found in the section (S4).

According to Bird et al. (2020), the generation time of the Réunion Harrier is 6.84 years. The 21-year study period therefore spans approximately three generations (20.52 years), the time window used by the IUCN Red List. From the trends computed over the three surveys (Eq.3), we estimated the yearly population growth rate *λ* over the 1998-2019 periods, i.e. 21 years, as well as the mean yearly change (see S9 & ?? ).

Models were fitted with R 4.3.2 (R Core Team, 2025) using the mgcv package (Wood, 2017). Following Wood (2011) we used the REML method, instead of the default Generalized Cross Validation (GCV) approach, to estimate smoothing parameters. Model fit was assessed visually, using the *mgcv* :: *gam*.*check*() and the *gratia* :: *appraise*() functions (see Fig. S6) and model residuals were also inspected to detect any problematic spatial pattern (spatial correlation of residuals was estimated with the *ncf* :: *sp*.*correlog*() function (Bjornstad, 2022), see Fig. S10 & Fig. S11). The dispersion parameter of the retained model based on Pearson residuals was 0.88; we did not consider this slight underdispersion problematic for parameters interpretation (Zuur *et al*., 2009). Details on model specifications and standard errors of different terms of the model presented in the results section care given in the sections S2, S7 and S8.

We used the packages *dplyr 1*.*1*.*4* (Wickham *et al*., 2022), *sf 1*.*0-21* (Pebesma & Bivand, 2023), *terra 1*.*8-60* (Hijmans, 2025), *tidyterra 0*.*7*.*2* (Hernangómez, 2023) and *openxlsx 4*.*2*.*8* (Schauberger & Walker, 2014) *to import and format the data, suncalc 0*.*5*.*1* (Thieurmel & Elmarhraoui, 2022) to compute sunrise schedule on La Réunion, the *gratia 0*.*11*.*1* package (Simpson, 2024) to assess and plot model fit of the GAM, and the following packages *ggplot2 4*.*0*.*0* (Wickham, 2016), *viridis 0*.*6*.*5* (Garnier *et al*., 2024), *ggpubr 0*.*6*.*1* (Kassambara, 2025), *ggridges 0*.*5*.*7* (Wilke, 2025), to produce the different figures.

Throughout the text, we report the mean values and the 95% confidence intervals.

## RESULTS

### Variations in abundance of breeding pairs in space and time

The retained model explained 17.6% of the deviance (adjusted *R*^2^ = 0.151). This model (Tab. 2) indicated that the number of breeding pairs recorded from the three surveys peaked in mid-June (Fig. 2a). Observation duration *Effort* had a positive effect on the number of pairs detected by observers: the longer the session, the more pairs were detected (Fig. 2b).

**Table 2:** Summary of the model retained to compute abundance index and population trends.

| **Component** | **Term** | **Estimate** | **Std Error** | **t-value** | **p-value** |  |
|---|---|---|---|---|---|---|
| A. parametric coefficients | (Intercept) | -0.58 | 0.06 | -9.24 | 0.00 | *** |
| **Component** | **Term** |  |  |  |  |  |
| B. smooth terms | s(yearF) | 2.14 | 6.00 | 0.60 | 0.12 |  |
|  | s(id_newF) | 0.00 | 344.00 | 0.00 | 0.62 |  |
|  | s(log_length) | 1.84 | 2.29 | 3.61 | 0.03 | * |
|  | s(jd_s) | 1.96 | 3.00 | 7.13 | 0.00 | *** |
|  | s(X,Y) | 24.52 | 34.41 | 1.14 | 0.26 |  |
|  | s(X,Y,surveyF) | 6.76 | 7.41 | 3.28 | 0.00 | ** |

Signif. codes: 0 <= '\*\*\*' < 0.001 < '\*\*' < 0.01 < '\*' < 0.05
Adjusted R-squared: 0.151, Deviance explained 0.176
-REML : 392.047, Scale est: 1.014, N: 994

**Table 3:** Annual population growth rate by area and period.

| Area | Period | Yearly *λ* | Yearly (%) |
|---|---|---|---|
| Island | 1998-2009 | 0.972 | -2.79 |
| East | 1998-2009 | 0.964 | -3.59 |
| West | 1998-2009 | 0.985 | -1.48 |
| Island | 1998-2017 | 0.978 | -2.20 |
| East | 1998-2017 | 0.973 | -2.70 |
| West | 1998-2017 | 0.986 | -1.39 |
| Island | 2009-2017 | 0.980 | -1.99 |
| East | 2009-2017 | 0.975 | -2.45 |
| West | 2009-2017 | 0.986 | -1.36 |

**Figure 2:**
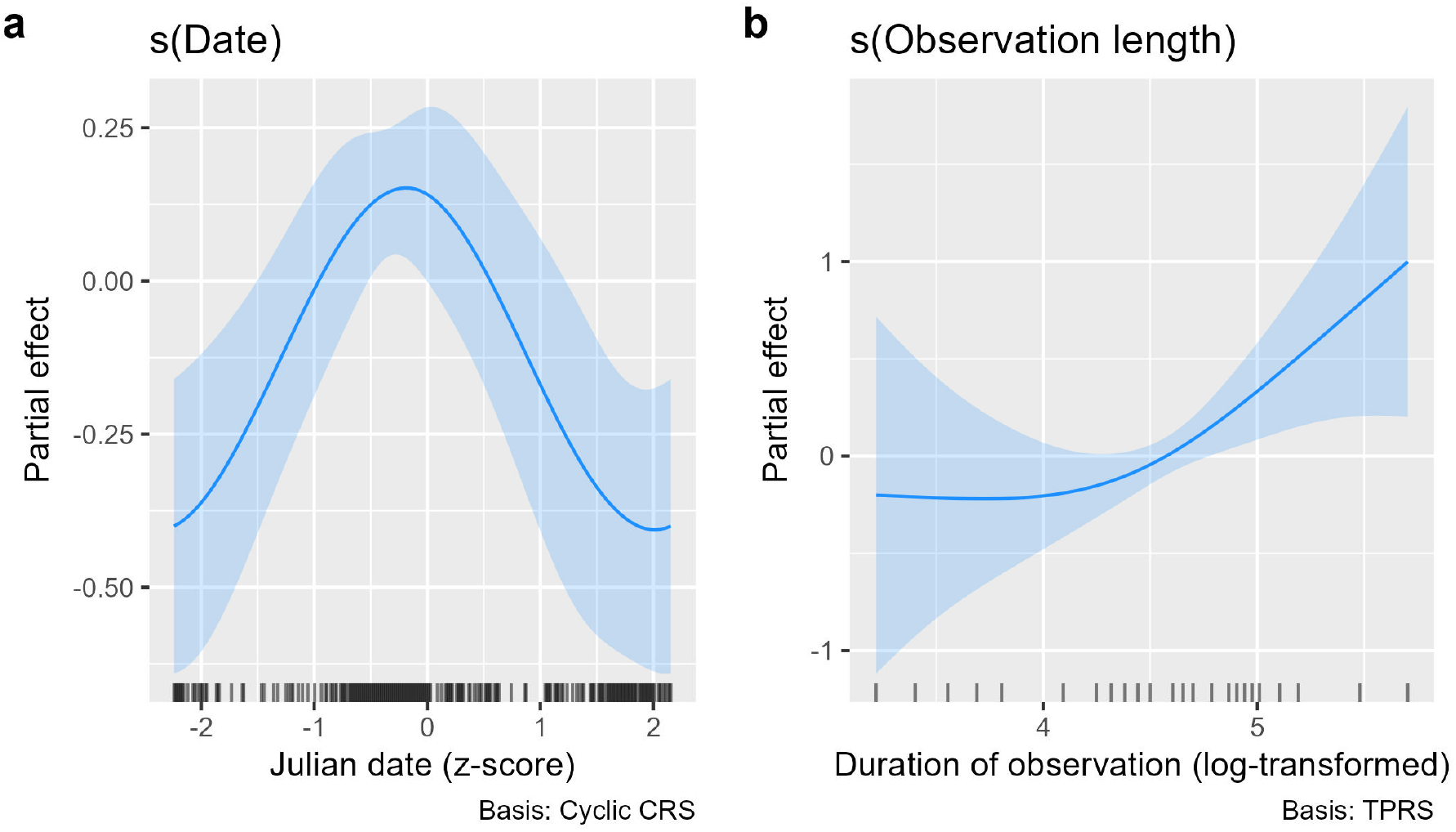
Relative effects on Julian date (a.) and duration of the observation (b.) from the selected model.

Areas of higher abundance, identified through the main spatial term *s*(*X, Y* ), were located mostly on the periphery of the island, at intermediate altitudes (Fig. 3a). Over the study period, the abundance index was particularly low around the Piton de la Fournaise, the active volcano of the island, and in densely populated areas of the western area of the island.

**Figure 3:**
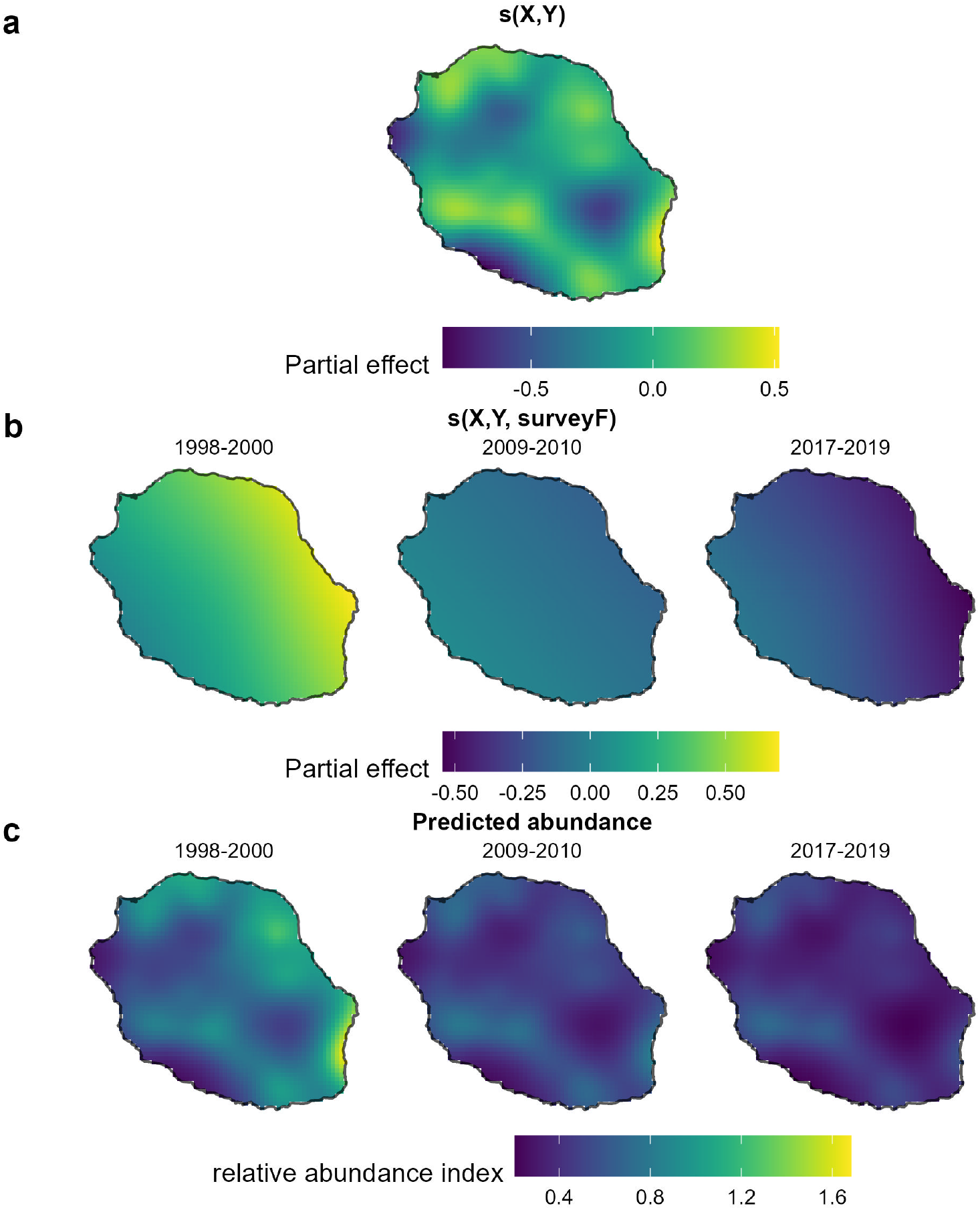
(a) Spatial variation in the relative abundance of Réunion Harrier across La Reunion, averaged across all 3 surveys, represented by the main spatial term s(X, Y ) in the model; (b) Deviation of each surveys from this average, given by the survey-specific term s(X, Y, surveyF ); (c) Predicted relative abundance for each survey, i.e. the sum of (a.) and (b.) plus an arbitrary constant.

The survey-specific terms *s*(*X, Y, surveyF* ), which capture how each survey’s spatial pattern departs from the main term *s*(*X, Y* ), indicate a complete shift in the distribution of abundance over the past 20 years (Fig. 3). The east-west gradient, initially higher in the east (1998-2000, Fig. 3b, left), was completely reversed by 2017-2019 (Fig. 3b, right). This is also visible in the predicted abundance (Fig. 3c), highlighting a more pronounced decline in the eastern part of the island than in the west.

### Population trends

When comparing changes in the abundance index at the island level (see Fig 4 a) & b), we found a strong decline between the 1998 and the 2009 surveys (-33.5% [-51.6%; -8.8%]), and a more uncertain, and likely weaker, decline between the 2009 and the 2017 survey with an estimated change of -17.9% [-42.5%; +14.4%]. Globally, the decline reached -46.2% [-61.7%;- 27.21%] between 1998 and 2017 surveys. Over the 12 years period separating the first two surveys, the population declined by an average of -2.78% per year (see SS9); over the whole study period, the yearly population change was -2.19%. Trends computed for the eastern and western parts of the island revealed a decline twice as high for the eastern part between the 1998 and 2009 (east: -43.1% [-61.8%; -17.1%] vs west -17.7% [-44.6%; +19.6%]) surveys or 1998 and 2017 surveys (east: -56.6% [-70.4%; -38.5%]) vs west: -29.6% [-53%; -0.3%]) between 1998 and 2017 surveys. For the eastern part of the island, this translates into an average annual decline of -3.59% and -2.69% over 12 and 21 years respectively, while for the west these values are lower, -1.47%. and -1.39%.

**Figure 4:**
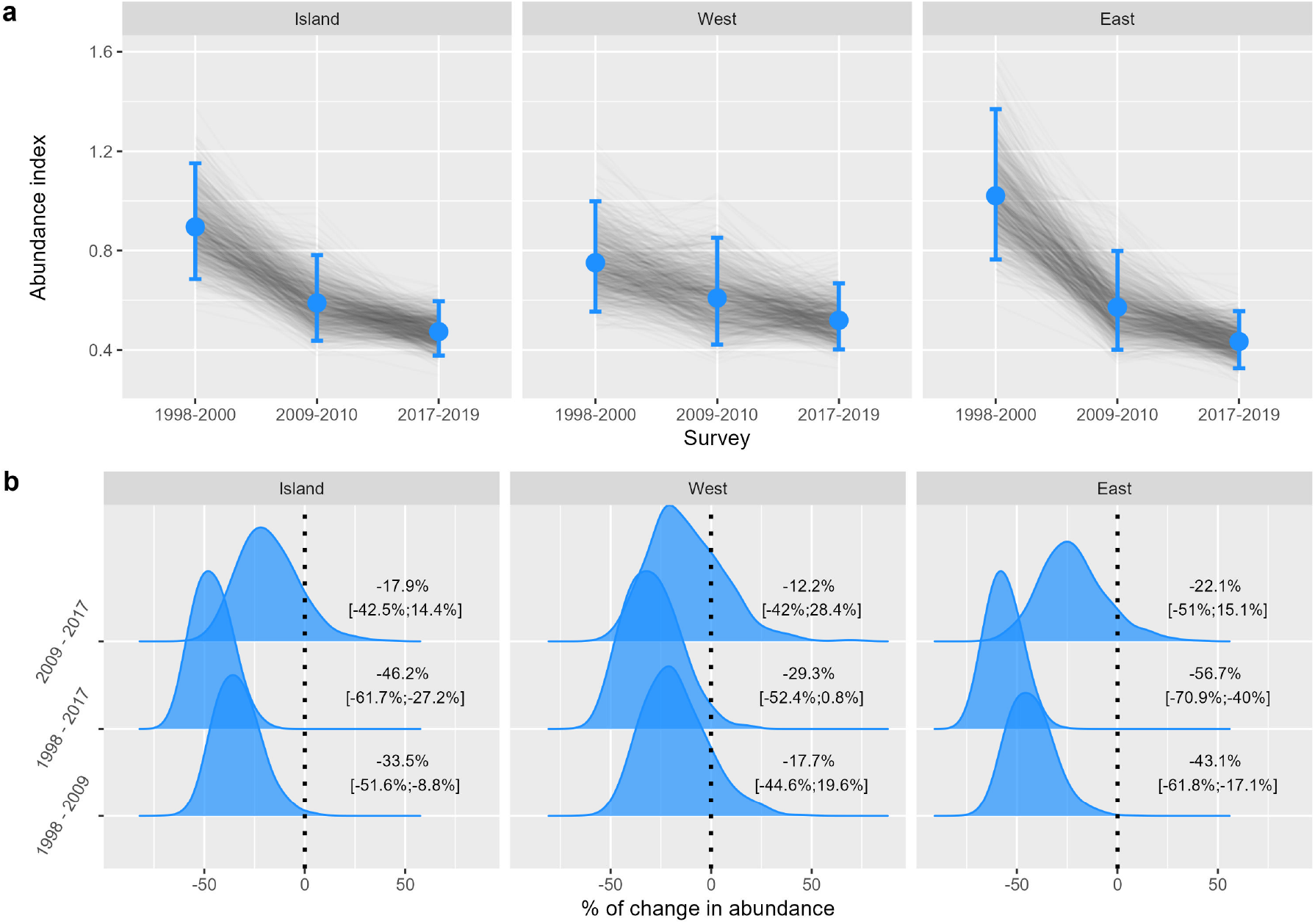
Predicted abundance index, averaged over 1,000 simulations, for the whole island (left) and for the western and eastern areas. Thin lines connect predicted values across surveys within a simulation; dots and error bars represent the mean and 95% interval of the 1,000 simulated values per survey. b. Distributions of predicted changes between pairs of surveys across simulations, with mean and confidence intervals shown to the right of each distribution. In (b), “1998”, “2009” and “2017” refer the 1998-2000, 2009-2010 and 2017-2019 surveys

## DISCUSSION

The present work fills an important gap in the conservation of the Réunion Harrier, the last bird of prey on this island, by providing the first objective estimate of the breeding population’s trend. The relative change in abundance was estimated to a mean -46% over the period 1998-2017, corresponding to three generations for this species. Using the “precautionary but realistic” approach recommended by IUCN, i.e. considering the lower bound of the confidence interval (-62%), the trend meets the -50% threshold of criterion A2b of the IUCN guidelines (IUCN, 2012), reinforcing evidence for the ‘Endangered’ (EN) status of *C. maillardi* (IUCN, 2012). Moreover, the relative abundance index displays contrasting trends, with a decline in the east twice as large as in the west (-56% vs -29%) between the two first surveys (1998-2009). Initially 25% higher in the East, the breeding index is now 15% lower than in the West, illustrating a marked spatial inversion. This suggests that the decline observed at the island level was mostly driven by a collapse of the breeding population of La Réunion Harrier in the eastern part of the island, with a more moderate decline observed in the western part. While the decrease is significant between 1998 and 2009, the decline is less pronounced, and non-significant, for the 2009-2017 period. It is however not possible to affirm that the population is now stable, given that the mean value of the trend for this period remains negative, which may indicate that detrimental factors persist. These results rely on three island-wide surveys spanning 21 years and on a statistical modelling approach that accounts for heterogeneity in dates and observation durations, as well as spatial heterogeneity in the sampling design. Our empirical evaluation of population dynamics is consistent with the demographic parameters reported by Fay et al. (2023). These authors showed that the species was characterised by low fecundity, resulting from small clutch sizes and a very low hatching success compared to other harrier or bird species in general, as well as a low adult survival for a bird of that size. While these values threaten the viability of the Réunion harrier, identifying the factors underlying these degraded parameters remains challenging, and the causal relationships have yet to be established.

Several non-exclusive pressures could contribute to these low demographic rates. In addition to genetic erosion and exposure to anticoagulant rodenticides, the Réunion Harrier is subject to direct persecution, collisions with power lines, wind turbines or vehicles, habitat loss and fragmentation, disturbance at breeding sites, variation in prey availability, predation, and extreme climatic events. However, the relative contribution of most of these pressures to the demography of the species remains poorly quantified. Available information currently suggests that illegal direct persecution accounts for relatively few recorded mortality events, although such events are inherently difficult to detect (Augiron, 2022). Collision risk and disturbance are increasingly considered through dedicated mitigation and spatial planning measures, but their population-level effects remain to be assessed. Therefore, the population decline documented here should probably be viewed as resulting from multiple, potentially interacting pressures rather than from a single identified driver. Among these pressures, recent work on genetic erosion and anticoagulant rodenticide exposure has received particular attention, as the available evidence allows their potential demographic consequences to be evaluated more directly.

Bourgeois et al. (2024) found that this species displays an overall very low genetic diversity, and one of the highest mutational loads ever recorded in a wild bird population, resulting from a severe bottleneck experienced over the last few millennia. During that period, the effective population size is estimated to have dropped from 3,000 to 300 individuals, with a sharper decline in the past century. In several vertebrate species, such erosion of genetic diversity has been shown to reduce individual fitness, with measurable effects on survival or breeding parameters (Kruuk *et al*., 2002; Huisman et al., 2016; Clarke et al., 2024), which can ultimately affect population trends (Blomqvist *et al*., 2010). The second factor is exposure to ARs which may negatively affect the species across the whole island. Since the late 1970s, these pesticides have been widely used to control rats in sugar cane fields, in other crops and around human habitations. Beyond their impact on agriculture (Stenseth *et al*., 2003), rats also carry spirochete bacteria that cause leptospirosis (Guernier *et al*., 2016), a zoonosis that can be fatal to humans (Allyn *et al*., 2024). Rats make up a significant portion of the Réunion Harrier’s diet, particularly in agricultural and urban areas (Augiron, 2022). Harriers can therefore become secondarily exposed to ARs, when feeding on contaminated rats. Coeurdassier *et al*. (2019) showed that rodenticides were very frequently encountered in harriers found dead, with 62% of the birds displaying a liver concentration in rodenticides known to be lethal in other raptors. Exposure to ARs, which increased between 1999 and 2015, may therefore contribute to additional mortality in both adults and juveniles. Although the effects of sub-lethal exposureto ARs are less well documented, they are suspected to reduce not only survival but also fecundity (Salim *et al*., 2014; Gomez et al., 2022).

One striking result of the present work is the differential population trends detected between the eastern and western regions of the island. This pattern mirrors the two genetic clusters identified by Bourgeois *et al*. (2024), located respectively in the western and eastern regions. While the species presents an overall low genetic diversity, the eastern sub-population exhibits an even lower level. These clusters may reflect restricted dispersal, in distance or frequency, arising from natural philopatry or from environmental constraints (landscape composition, natural barriers) that limit movements, even in flying vertebrates such as bats (Pinaud *et al*., 2018) or birds (Martensen *et al*., 2008). For example, a small passerine, the Réunion Grey White-eye *Zosterops borbonicus*, is characterised by an exceptionally high level of intra-island genetic differentiation that appears to be mostly shaped by restricted dispersal, and to a lesser extent landscape composition (Milá *et al*., 2010; Bertrand et al., 2014). Very few dispersal events have been documented in the Réunion Harrier so far (Augiron, 2022), whether with wing-tagged individuals or satellite tracking. It remains however unclear whether this limited dispersal is due to habitat features, e.g. the volcanic range dividing the island, or to the expansion of anthropogenic habitats since human settlement. Approximately 50% of the island is now covered by cultivated areas, urban areas, or secondary vegetation (Strasberg *et al*., 2005), which may have further disrupted dispersal processes. The loss of key landscape elements, together with the presence of built-up areas, may limit the ability of birds to move across this altered landscape, thereby reinforcing the spatial isolation of genetic clusters.

No spatial data on rodenticide use are available for the eastern and western parts of the island, so we cannot assess whether it explains the contrasting trends we detected. However Coeurdassier et al. (2019) found a heterogeneous spatial pattern for exposed individuals, with most exposed birds found in the east. Assuming that the probability of discovering a dead or injured harrier is spatially homogeneous, this would suggest birds are indeed more exposed in this part of the island. A plausible but speculative explanation for this higher exposure risk could lie in rodents availability. indeed, rainfall is known to increase rat abundance in tropical regions (Madsen & Shine, 1999), and the two areas have strikingly different regimes (2,132 mm/year in the east vs 886 mm/year in the west). The east may therefore offer a greater availability of rats, exposing harriers to a higher risk of lethal and sub-lethal effects. Taken together, genetic erosion and rodenticide exposure are likely to interact, creating a self-reinforcing process typical of extinction vortices (Fagan & Holmes, 2006). High mortality reduces population size and dispersal opportunities, which limits genetic diversity; this in turn further decreases fitness and exacerbates demographic decline.

To halt the decline and prevent the species from approaching extinction, the impact of the pressures acting on the population must be substantially reduced without delay. Given the species’ conservation status, complementary conservation actions should be implemented simultaneously, while continuing to improve knowledge of its ecology and demography. Among currently documented threats, reducing harriers’ exposure to anticoagulant rodenticides would have a strong positive effect on population dynamics by lowering adult and juvenile mortality, and potentially improving breeding success (Coeurdassier *et al*., 2019; Salim et al., 2014; Gomez *et al*., 2022). Under the European legal framework governing the renewal of plant protection product authorisations (Regulation (EC) No 1107/2009), the ongoing RatEx pilot study aims to reduce ARs’ use by identifying areas where is unnecessary, either because leptospirosis is not circulating locally or because rat densities do threaten crops. However, reducing ARs use while protecting public health and crop yields will likely require years of cross-sectoral effort. In parallel, other conservation measures should be deployed or reinforced, including reducing collision risks with power lines, wind turbines and vehicles (Bellebaum *et al*., 2013; Slater et al., 2020; Bullock et al., 2024), protecting breeding and foraging habitats, maintaining landscape connectivity (Strasberg *et al*., 2005), and limiting disturbance at key breeding sites. Such actions should be spatially prioritised according to breeding abundance, habitat suitability and identified risk areas.

Given the very low genetic diversity, high mutational load and limited dispersal documented for the species (Bourgeois *et al*., 2024; Augiron, 2022), genetic management could also represent a complementary conservation option. Genetic rescue has proven to be efficient in several conservation programs worldwide (Bouzat *et al*., 2009; Heber et al., 2013; Wright et al., 2014; Lewanski *et al*., 2025), although the species-specific ecology and demographic constraints must be carefully considered (Nichols *et al*., 2024). For the Réunion Harrier, potential approaches could include translocation of eggs or chicks between eastern and western breeding sectors (Martínez-Cruz, 2011), possibly combined with temporary captive rearing and release. Another option could involve collecting one egg per clutch from selected wild nests for artificial incubation and captive rearing at a wildlife rescue centre, followed by release into the wild. The latter might, however, prove challenging since hatching success is currently as low as 40% (Fay *et al*., 2023), reducing the chance that a collected egg would actually be viable. Finally, a captive breeding program should only be considered as a long-term, last-resort option (Heinrichs *et al*., 2019; Witzenberger & Hochkirch, 2011), as harriers are especially difficult to keep in captivity. Importantly, such genetic interventions should complement, rather than replace, actions aimed at reducing the underlying causes of mortality and reproductive failure.

The spatial abundance index produced in this work highlights hotspots for the species. The last survey (2017–2019) was designed to allow the explicit estimation of detection probability and total population size, including the size of the breeding population (Couturier *et al*., 2022, & work in progress). Combined with information on suitable breeding habitats and spatially explicit threats, this information can help identifying priority conservation areas, targeting management actions across the island, and improving the integration of harriers conservation into spatial planning and forthcoming prefectural regulations. In particular, it could help identifying areas where rodenticide reduction, mitigation of collision risks or disturbance limitation should be prioritised.

Establishing a new study site in the western region would provide valuable data from a distinct ecological context, help to identify the factors driving population dynamics (Robinson *et al*., 2014), allowing a more robust assessment of threats, yield more accurate predictions of population trajectories, and support the design of adaptive conservation strategies (Wiens *et al*., 2017; Winship et al., 2013). The new standardised protocol, relying heavily on citizen participation, is an invaluable tool for monitoring future population changes and assessing the effectiveness of conservation actions. It addresses needs identified in the National Species Action Plan (2022-2031), to which the present work directly contributes.

## ACKNOWLEDGMENTS

We sincerely thank all the observers, volunteers, and professionals, who took part in the surveys over the past 20 years; their efforts made it possible to build a dataset supporting a temporal indicator of the breeding population, essential for the conservation of this endangered species.

We are grateful to the members of the Société d’Etudes Ornithologiques de La Réunion for organizing simultaneous harrier counts across the entire island. We also extend our thanks to the many funding partners who supported this long-term monitoring programme over the past two decades: the Regional Council of the Réunion (grant), the European Union (FEDER POT and FEDER Grant n°RE0005162), the French Ministry of Environment (DIREN-Réunion), the DEAL Réunion, Aerowatts, the Etang-Salé municipality, EDF and Tereos. The authors are grateful to Drs. C. Morrison and R. Simmons, as well as two anonymous reviewers, for their thoughtful comments, and to Pr. N. Yoccoz, who acted as *PCI* recommender.

## Authors’ contribution

- Alexandre Villers: Protocol design, Supervision, Data curation, Analytical methodology, Formal analysis, Writing - original draft.
- Thibaut Couturier: Protocol design, Project management and administration (2017), Data curation, Field data sampling, Reviewing and Editing
- Pierrick Ferret: Protocol design, Data curation, Field data sampling, Reviewing and Editing
- Damien Chiron: Protocol design, Project management and administration (2018-2019), Data curation, Field data sampling, Reviewing and Editing
- Nicolas Laurent: Protocol design, Field data sampling, Design of web interface and data management, Reviewing and Editing
- Thomas Cornulier: Protocol design, Analytical methodology, Formal analysis, Reviewing and Editing
- Aurélien Besnard: Protocol design, Analytical methodology, Reviewing and Editing
- Vincent Bretagnolle: Field data sampling, Data curation, Reviewing and Editing
- Marc Salamolard: Project management and administration (1998-2000), Field data sampling, Reviewing and Editing
- Valérie Grondin: Project management and administration (2009-2010), Field data sampling, Data curation, Reviewing and Editing
- François-Xavier Couzi: Field data sampling, Project Administration, Reviewing and Editing
- Rémi Fay: Analytical methodology, Reviewing and Editing
- Cyril Eraud: Analytical methodology, Reviewing and Editing
- Steve Augiron: Protocol design, Funding acquisition and management, Project management and administration (2016-2019), Field data sampling, Supervision, Data curation, Formal analysis, Writing - original draft.

## ADDITIONAL INFORMATION

### Access to data and code

The data and code supporting this work are archived on Zenodo DOI: 10.5281/zenodo.24374859 and released under a CC BY 4.0 license.

### Use of Artificial Intelligence

after the first round of reviewing, the consolidated version of the manuscript was submitted to Chat GPT-5.6 Luna and Claude Sonnet 5.5 with specific directives to review English grammar and formulations. None of the analytical framework nor coding was produced with AI.

## Supplementary material

### S1 Distribution of sampling effort in space and time

Distribution of sampling points monitored for each survey, for the whole data set (upper panels), during the main breeding period, from April to June (central panels) or outside breeding (lower panels).

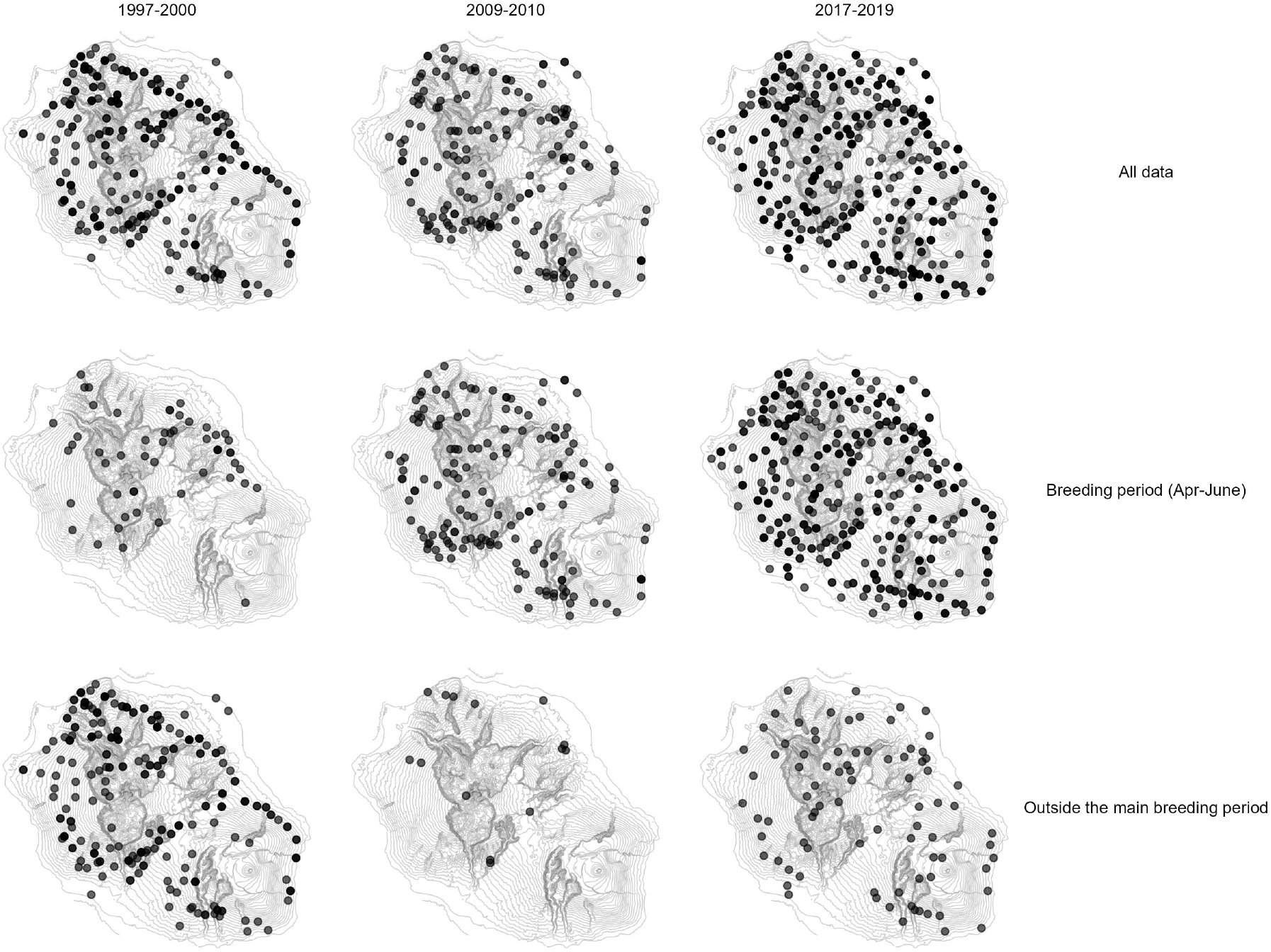

### S2 Details on GAM specifications

The date of the observation session was expressed as a Julian date variable “Date”, with 1^st^ of January corresponding to day 1. This term was specified with a cyclic cubic regression spline, in order to account for the fact that the 31^*st*^ of December and the 1^*st*^ of January are equivalent from a biological point of view, e.g. in term of the phase of the breeding cycle or expected climatic conditions at this time of the year. The value for the dimension of the basis function was set to *k* = 5 to match *a priori* expectations and to limit the risk of overfitting. The length of the observation session (log-transformed), “Effort”, was modelled with a thin plate regression spline with dimension also set to a maximum of *k* = 5. Spatial smoothers i.e. an interaction between latitude and longitude, were also included in *f*_3_ and *f*_4_ to model potential spatial autocorrelation. These terms account for local specificities (habitat, abiotic conditions) and for heterogeneity in the sampling design, notably the wider range of environments covered in the last survey (parametrisation in S3).

The retained model included the main interaction between latitude and longitude, as well as smooth terms modelling deviations from the main spatial smooth for each survey *SurveyF* . Two random intercepts were also included, namely *Y earF*, a 9-level random effect accounting for yearly variations of abundance, and *SiteID*, a 346-level random effect accounting for the identity of sampling points, with identity shared between and within surveys to account for inter- and intra-annual repeated counts. We assumed a Tweedie distribution (log link) a family that encompasses continuous (normal, gamma, inverse Gaussian), discrete (Poisson) and compound (Poisson-gamma) distributions. This distribution has three parameters, the mean (*µ*), a dispersion parameter (*ϕ*) and a power parameter (*p*). In the *mgcv* package, the value of *p* is restricted to ]1; 2[, and is estimated simultaneously with other parameters, offering flexibility in the shape of the probability distribution assumed for the response, particularly with respect to overdispersion and excess zeros, when needed.

### S3 Parametrisation of the spatial term and model specifications

We set the value of the basis dimension of the spatial term to *k* = 100, in order to ensure that a sufficient number of degrees of freedom would be available for this smoother. We compared four specifications of the spatial term and retained the one that, in our judgement, fitted best, since Generalized Additive Modelling often relies on expert knowledge (Pedersen *et al*., 2019). AIC-based selection,although common, performed poorly given the complexity of spatial terms and spatio-temporal interaction formulations . The first model included a *surveyF* fixed effect factor and a spatial smoother common to the whole period of survey with the term *s*(*X, Y, k* = 100). Two models included a smooth factor interaction between coordinates and the survey period *s*(*X, Y, k* = 100) +*s*(*X, Y, SurveyF, bs* = “*sz*”, *k* = 100) or *surveyF* +*s*(*X, Y, k* = 100) + *s*(*X, Y, by* = *SurveyF, m* = 1, *k* = 100). The latter models deviations from the main spatial term for each survey (a 3-level factor), with independent smoothness / wiggliness for each level. With *m* = 1, the marginal thin-plate spline basis is penalised on the squared first derivative of the function, rather than the second, which limits collinearity between the global smoother *s*(*X, Y*) and the survey-specific terms *s*(*X, Y, by* = *SurveyF*).

The data and code supporting this work are archived on Zenodo (see

### S4 Computing confidence intervals for trends

We combined predictions and simulations to obtain the distribution of the abundance index, thereby propagating parameter uncertainty. First, using the *lpmatrix* option in the *predict*.*gam*() function, we generated a model matrix *X*_*p*_ for prediction. Values of explanatory variables for computing *X*_*p*_ were set as follows: coordinates were those of the centroid of each 1km x 1km grid cell, Date and Effort were set at their respective means, random effects *Y earF* and *SiteID* were set to a fixed arbitrary level of each factor. This setup was replicated for all surveys (indicated by column *SurveyF* ), to generate predictions of relative abundance for each survey. Second, we generated a *B*_*r*_ matrix where each line was a set of model parameters, corresponding to columns of *X*_*p*_, and each column was a replicate parameter vector. Parameter replicates (n=1000) were simulated from their posterior distribution using the mgcv::rmvn() function, with the variance-covariance matrix estimated with the *mgcv* :: *vcov*() function. Finally, the matrix *P*, containing for each grid cell and survey, 1000 sets of simulated abundance values, was computed such that:

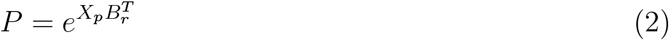

Trends were expressed as:

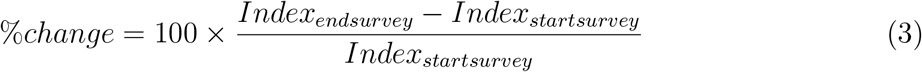

To estimate population trends between surveys at the island level, we averaged, for each replicate, the value of the abundance index for all grid cells covering the island. We then compared the trends in population between pairs of surveys following Eq.(3). To represent spatial changes in trends between surveys, we took, for each pair of surveys and for each grid cell, the 2.5th and 97.5th percentiles of the trend distribution *stats* :: *quantile*() as 95U% confidence intervals.

### S5 Effect of data selection on trends

We fitted a model of the form

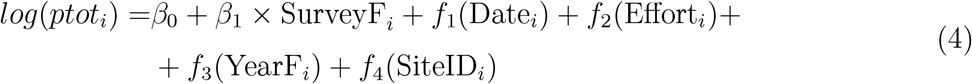

to 4 different data sets: the full data set; only the points common to the three surveys; only the points surveyed between April and June; and the points common the three surveys and surveyed between April and June.

Trends computed at the island level are compared in the figure below.

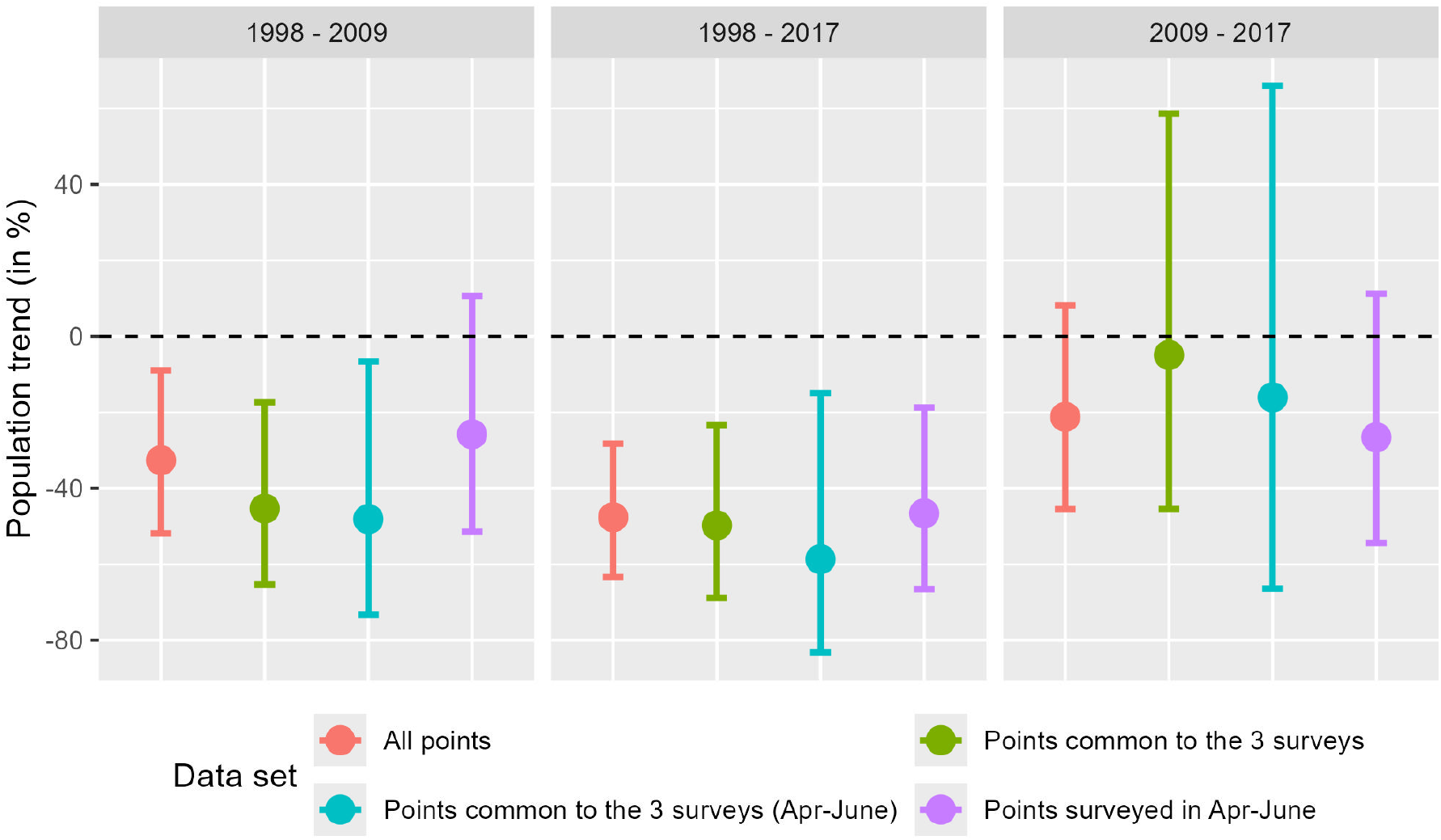

### S6 Assessment of best model fit

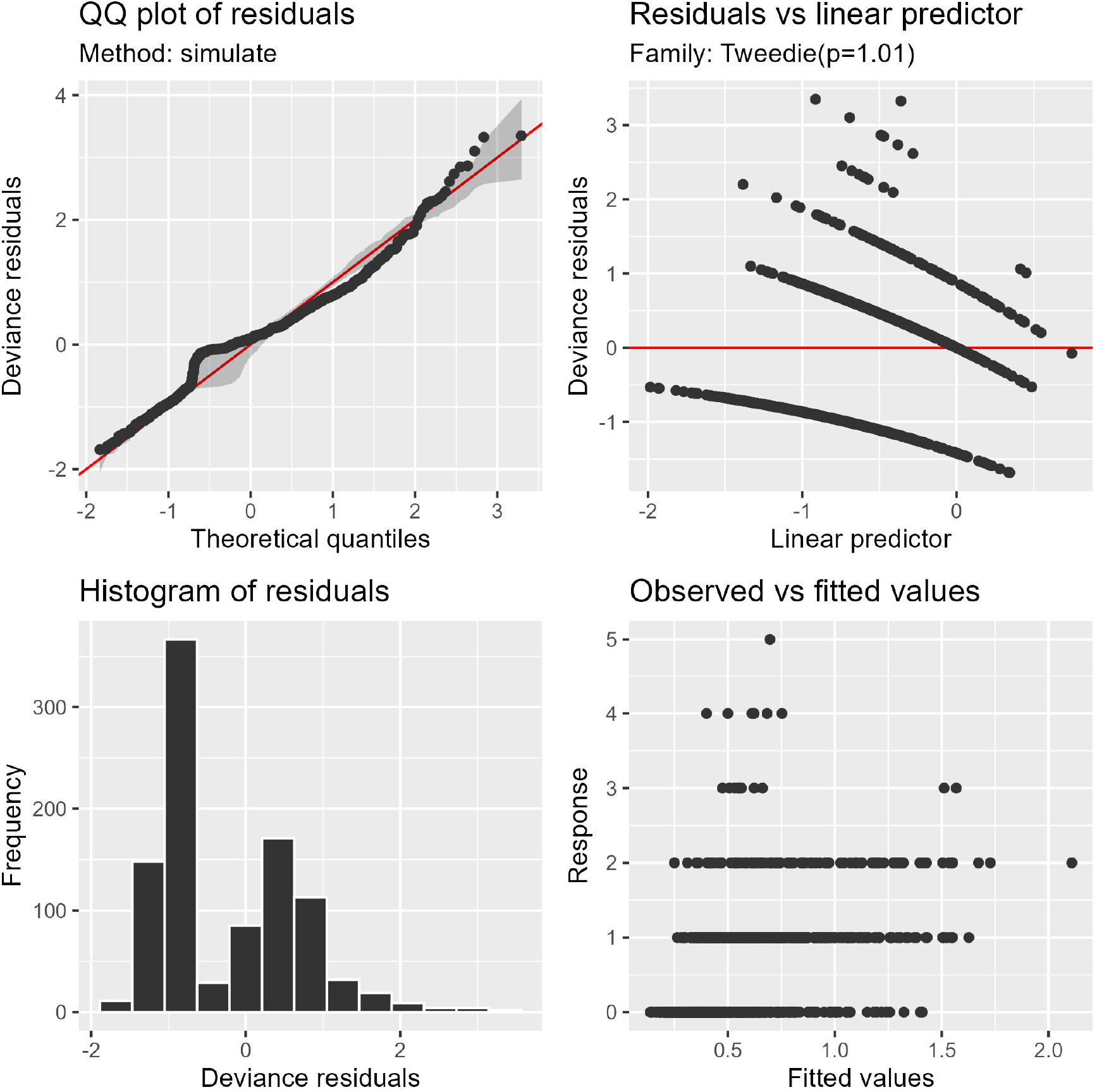

### S7 Spatial terms of the GAM: means and standard error

a. Spatial effect modelled across the 3 surveys represented by the spatial smoother *s*(*X, Y* ); and b. the associated standard error; c. Survey specific spatial smoother *s*(*X, Y, surveyF* ), modelling the spatialised difference in abundance between the main spatial smoother *s*(*X, Y* ) and each survey; and d. the associated standard error of this term across the three surveys.

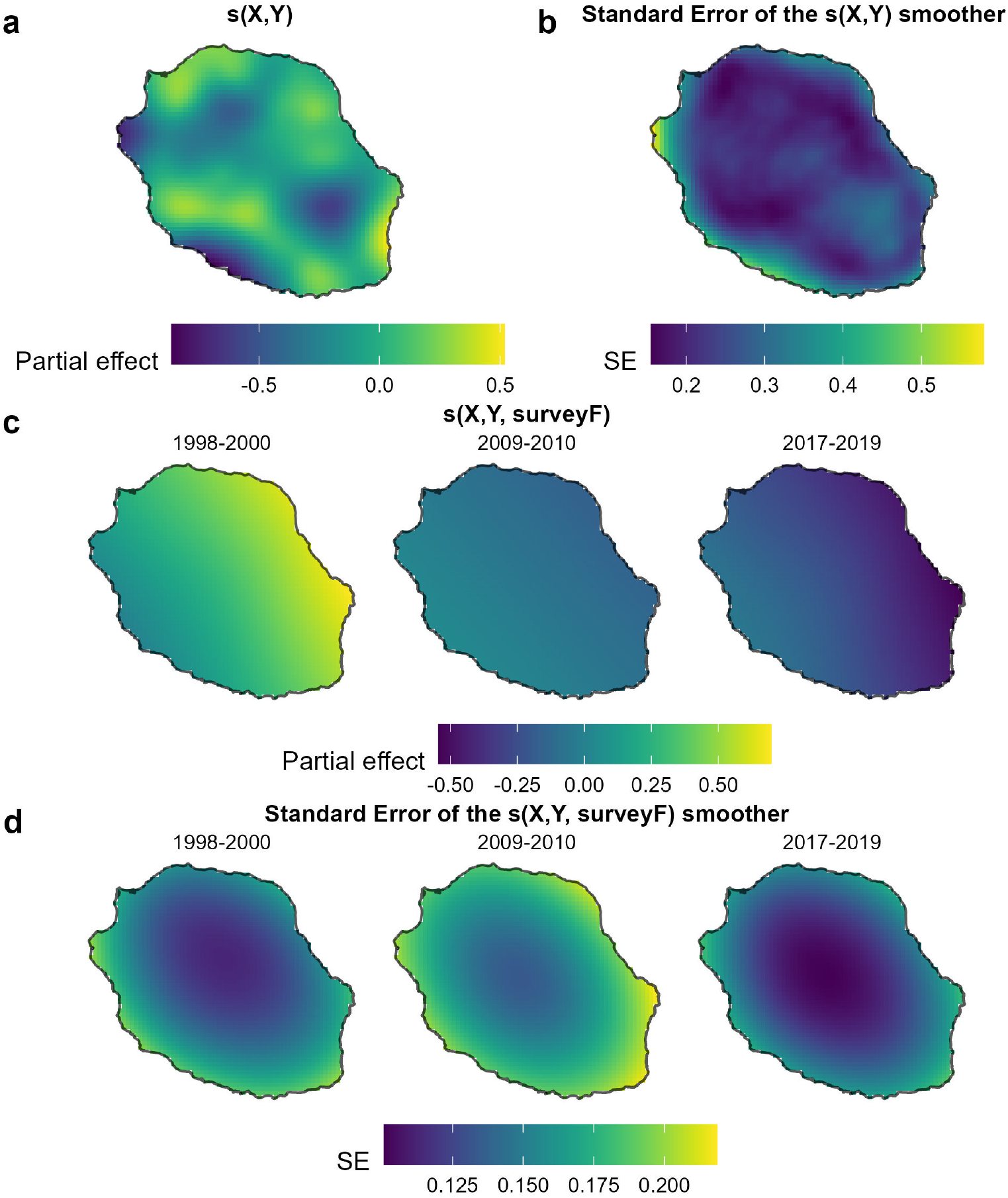

### S8 Predicted abundance in space: means and standard error

a. Predicted abundance index for the three surveys; and b. associated standard error of the mean abundance. Red dots indicate the spatial distribution of the sampling points of each survey.

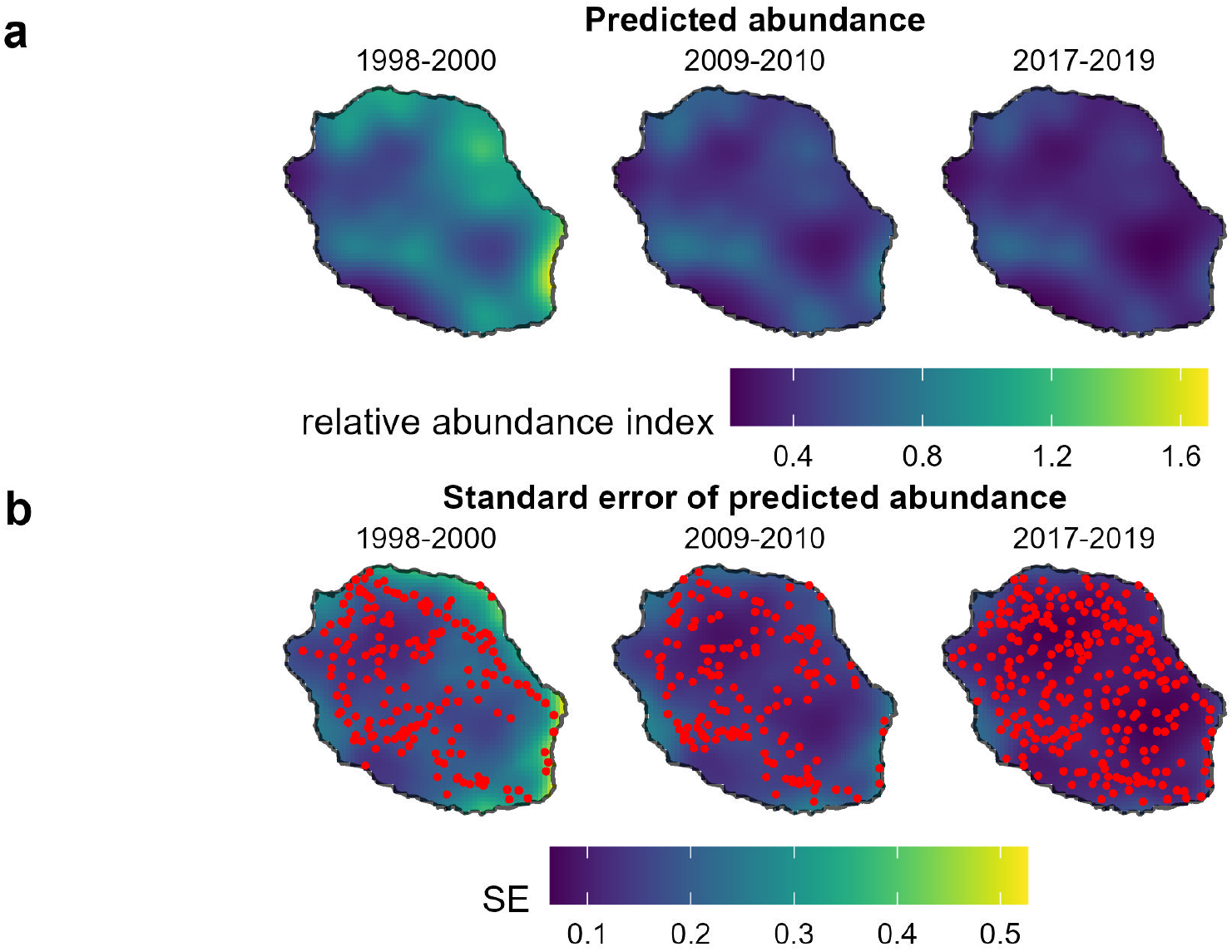

### S9 Population growth rate and yearly rate of change

### S10 Spatialised residuals

Residuals values for all the data and each of the three surveys

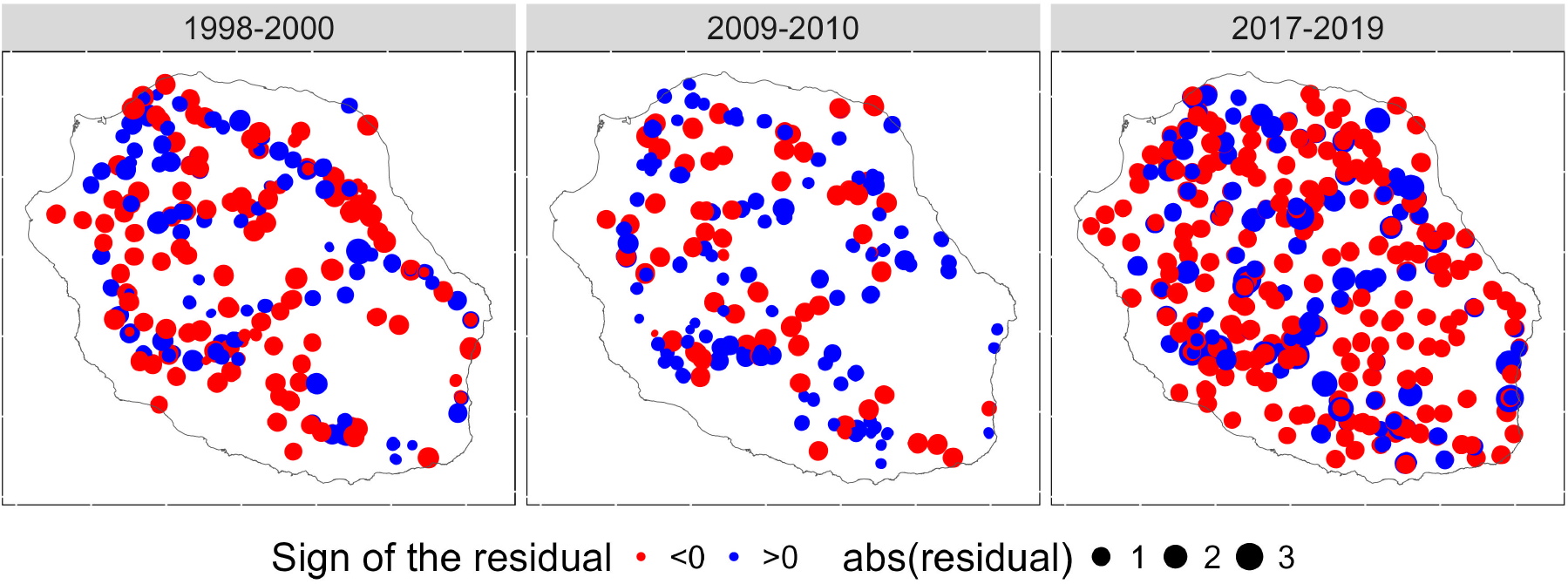

### S11 Spatial correlograms of residuals of the selected model

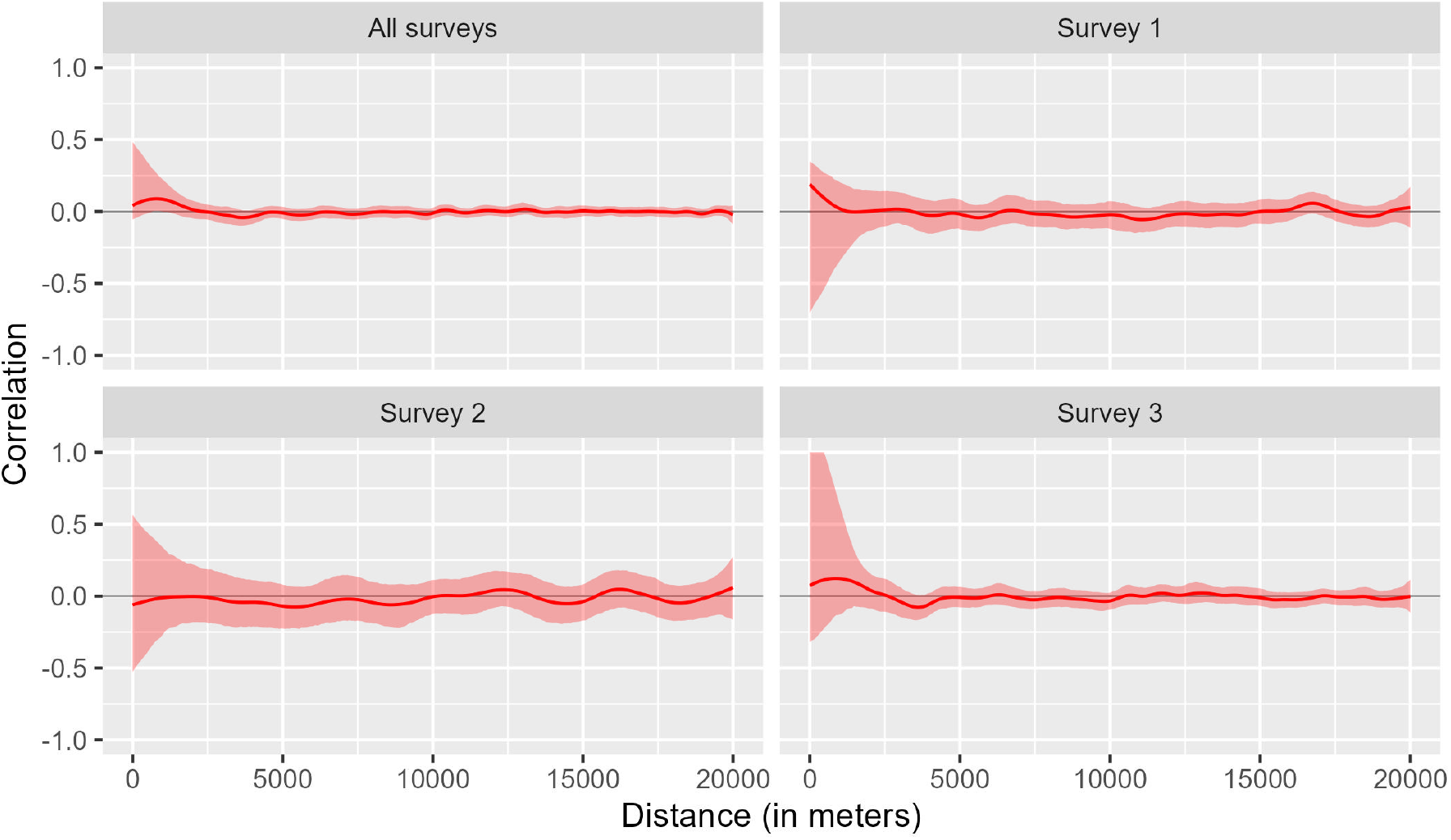

